# Phased genome assemblies and pangenome graphs of human populations of Japan and Saudi Arabia

**DOI:** 10.1101/2024.12.18.628902

**Authors:** Maxat Kulmanov, Saeideh Ashouri, Yang Liu, Marwa Abdelhakim, Ebtehal Alsolme, Masao Nagasaki, Yasuyuki Ohkawa, Yutaka Suzuki, Rund Tawfiq, Katsushi Tokunaga, Toshiaki Katayama, Malak S Abedalthagafi, Robert Hoehndorf, Yosuke Kawai

**Author notes:** Corresponding authors Correspondence to Malak S. Abedalthagafi, Robert Hoehndorf, and Yosuke Kawai;. These authors contributed equally (Maxat Kulmanov, Saeideh Ashouri). These authors contributed equally (Malak S Abedalthagafi, Robert Hoehndorf, Yosuke Kawai).

## Abstract

The selection of a reference sequence in genome analysis is critical, as it serves as the foundation for all downstream analyses. Recently, the pangenome graph has been proposed as a data model that incorporates haplotypes from multiple individuals. Here we present JaSaPaGe, a pangenome graph reference for Saudi Arabian and Japanese populations, both of which have been significantly underrepresented in previous genomic studies. We constructed JaSaPaGe from high-quality phased diploid assemblies which were made utilizing PacBio high-fidelity long reads, Nanopore long reads, and Hi-C short reads of 9 Saudi and 10 Japanese individuals. Quality evaluation of the pangenome graph by variant calling showed that our pangenome outperformed earlier linear reference genomes (GRCh38 and T2T-CHM13) and showed comparable performance to the pangenome graph provided by the Human Pangenome Reference Consortium (HPRC), with more variants found in Japanese and Saudi samples using their population-specific pangenomes. This pangenome reference will serve as a valuable resource for both the research and clinical communities in Japan and Saudi Arabia.

## Background & Summary

Pangenomes are the set of all genes in a species or population, and pangenome graphs are data structures to represent pangenomes. Recently, pangenome graphs were made available to be used as reference for human genomics instead of linear reference genomes [1]. Pangenome graphs have the advantage that they include multiple genomes and, consequently, more variation than single genomes. In particular, they include additional structural variants, providing a more flexible and inclusive framework for aligning reads, resulting in increased alignment rates and reduced mapping errors, and, therefore, identifying genomic variants, including potentially disease-causing variants [2]. Pangenome graphs are not only valuable data structures for representing multiple genomes, but their application can also significantly enhance the success of genomics and precision medicine.

Genetic variation is population-specific. Although the draft human reference pangenome graph includes individuals from diverse regions [1], it significantly lacks representation from Arab and Japanese populations, collectively accounting for approximately 600 million people worldwide.

Moreover, the question of how many individuals from a population should be included in a pangenome graph to adequately capture sufficient variation for accurate variant calling remains unresolved. Ideally, the graph should capture most of the genetic variation, such that adding additional individuals to the reference does not significantly improve the performance of bioinformatics workflows relying on it. The optimal number of individuals depends on their specific genotypes, population structure, and genetic diversity within the population.

While it has become technically possible to assemble human genomes *de novo* due to advancements in long-read sequencing technology, only few human genomes have been made available freely without restrictions, as would be required for a reference genome or pangenome. Current limitations include the limited number and quality of publicly available genomes, technical limitations related to the scale and complexity of pangenome graphs and their software, and ethical concerns, as only a relatively small number of human samples have sufficient ethical approval for unrestricted research use [3].

Pangenome graphs are relatively new types of data structures, and the technology surrounding them is rapidly evolving and still maturing [4]. While bioinformatics tools and methods have evolved a long time around single and linear reference genomes, pangenome graphs still face some limitations in available algorithms and software [5]. Consequently, the pangenome graphs cannot yet grow to arbitrary sizes including thousands of individuals from many different populations, and they may never grow to these sizes due to general resource limitations, particularly in regions of the world where computational resources are not easily available. Therefore, population-specific pangenome graphs will likely play a prominent role in population-specific genomic analysis [6], [7], including in clinical genetics, and several population-specific pangenome graphs are now becoming available [8] [9].

We utilized multiple sequencing technologies to sequence 10 Japanese and 8 Saudi individuals, creating phased diploid assemblies for each sample. These assemblies were used to develop two population-specific pangenome graphs: a Japanese pangenome graph derived from 10 Japanese individuals, and a Saudi pangenome graph generated from 8 Saudi individuals combined with one publicly available Saudi diploid genome [10], totaling 9 Saudi samples.

We further combined the two population-specific pangenome graphs into a panpopulation pangenome graph, consisting of two genetically distinct populations and two common reference genomes (T2T-CHM13 and GRCh38). The sequencing data, the genome assemblies, and the pangenome graphs, together with the software and scripts used to generate and evaluate them, are freely available following the FAIR (Findable, Accessible, Interoperable, Reusable) principles [11].

## Methods

### Sample processing and sequencing

### Saudi samples

Fresh blood was collected from the donors and shipped to KAUST (Thuwal, Saudi Arabia) in ambient temperature for DNA extraction. All samples arrived at KAUST at most 24 hours after blood was taken from the donors. DNA was extracted using two methods. Ultra-high molecular weight DNA (uHMW) was isolated from fresh blood samples using New England Biolabs (NEB) Monarch High Molecular Weight (HMW) DNA isolation kit following the manufacturer’s protocol (New England Biolabs, UK) with a minor modification: agitation was set at 700 rpm during the lysis step. DNA was kept at 4°C until library preparation for long-read sequencing using the PacBio and Oxford nanopore sequencing platforms. The second DNA extraction method was the initial step of the Dovetail Omni-C protocol, employed for constructing Hi-C libraries following the manufacturer’s protocol.

We established criteria for sample selection that balanced quality requirements with practical constraints. While the recommended requirements call for 30-40 μg of total DNA with an A260/A230 ratio greater than 2.0, we adjusted our thresholds based on sample availability and project needs. Our minimum criteria included a total DNA amount of at least 13 μg (the lowest we processed), a concentration not less than 40 ng/μL, an A260/A230 ratio not below 1.9, an A260/A280 ratio between 1.8 and 2.0, and an average fragment size exceeding 100 kb. We strictly adhered to the 40 ng/μL concentration threshold to ensure sufficient DNA density for optimal sequencing performance. These adjusted parameters allowed us to include samples that would have been excluded under strict adherence to ideal standards while maintaining a minimum quality baseline. Samples meeting these criteria were chosen for sequencing on ONT.

We used three platforms for sequencing the extracted DNA: Illumina NovaSeq 6000 (Illumina, San Diego, USA), PacBio Sequel II (Pacific Biosciences, Menlo Park, USA), and Oxford Nanopore PromethION (Oxford Nanopore Technologies, Oxford, UK). We employed the Dovetail Omni-C protocol to construct Hi-C libraries (Dovetail Genomics, USA). The Dovetail Omni-C protocol is a widely adopted method that allows for the interrogation of chromatin interactions on a genome-wide scale. This protocol incorporates a series of steps to capture spatial proximity information between genomic loci, enabling the investigation of three-dimensional chromatin organization. In brief, the protocol involves crosslinking and restriction enzyme digestion of chromatin, followed by ligation of biotinylated adapters to the digested fragments. The ligated DNA is then sheared, and proximity ligation is performed to capture chromatin interactions. Each resulting DNA library was sequenced on one lane of an SP flowcell to enable the identification and characterization of chromatin interactions at high resolution. High molecular weight gDNA was sheared with Megaruptor 3 (Diagenode, Denville, USA) to the size range of 15-20 kb. SMRTbell was prepared with HiFi Express Template prep kit 2.0 (102-088-900), and size-selected with the PippinHT System (Sage Science HTP0001). Finally, SMRTbell QC was assessed with Qubit dsDNA High Sensitivity (model Q33230; ThermoFisher Scientific, Waltham, USA) and FEMTO Pulse (Inc. P-0003-0817; Agilent Technologies, Santa Clara, USA). Sequencing of SMRTbell was set up on PacBio Sequel II system with Sequel II Binding kit 2.2 (101-894-200), Sequel II Sequencing Kit 2.0 (101-820-200), and SMRTcell 8M Tray (101-389-001), according to conditions specified in SMRTlink with 30 hour movie times, 2 hour pre-extension time, and adaptive loading mode.

For Nanopore sequencing, for each sample, two libraries were prepared using the Ultra-Long Sequencing Kit (SQK-ULK001) and its recommended protocol from Oxford Nanopore Technologies (Oxford Nanopore Technologies, Oxford, UK). 30 μg of uHMW gDNA was used as input for each library. A PromethION flow cell (FLO-PRO002) was primed and prepared according to the same protocol, and 75 μl of a sequencing library was loaded. Twenty-four hours after the first sequencing run, a nuclease flush and priming step was performed according to the protocol, and an additional 75 μl of library was loaded before starting another sequencing run. This process was repeated until each library was loaded three times. Basecalling was performed using Guppy v5.1.13 with the super-accurate basecalling model. The process was repeated with the second prepared library and another PromethION flow cell.

### Japanese samples

We selected ten male samples of JPT (Japanese in Tokyo, Japan) from the HapMap collection [12]. We obtained LCL cell lines from the NHGRI Sample Repository for Human Genetic Research at the Coriell Institute for Medical Research, and extracted the DNA from these cell lines. In some analyses, we used extracted DNA provided by the repository. DNA was extracted from cultivated cells using the Nanobind CBB Big DNA Kit (Pacific Biosciences, Menlo Park, USA) following the manufacturer’s protocol. The extracted DNA samples were delivered for long-read sequencing using the PacBio and Oxford nanopore sequencing platforms.

For the PacBio sequencing, the size distribution of gDNA was assessed using a FEMTO Pulse, and the concentration was measured with Qubit 3.0 using the dsDNA BR and RNA BR assay kits, as well as with a Nanodrop (ThermoFisher Scientific, Waltham, USA). After confirming that the gDNA met PacBio’s recommended quality standards, gDNA was fragmented. We used 5 μg of gDNA and fragmented it to an average size of 16–20 Kbp using a Megaruptor 3 and the Megaruptor Shearing Kit/DNAFluid+. Library preparation was performed using the SMRTbell prep kit 3.0, following the manufacturer’s protocol. For final library size selection, a PippinHT System with 0.75% agarose gel cassettes and marker S1 was used. The cut-off size range was set to 10 Kbp–50 Kbp as recommended by the manufacturer. Subsequently, library QC was performed using FEMTO Pulse and Qubit 1x dsDNA HS kit. The prepared sequencing templates were loaded onto the Sequel IIe system using Binding kit 3.2 and cleanup beads (>3 Kbp). A 30-hour movie run was performed, generating high-fidelity (Hi-Fi) reads.

Nanopore sequencing was performed using the conventional long-read sequencing that had been previously analyzed, with additional ultra-long read sequencing performed on four samples. The sequencing protocols used were the Ligation Sequencing Kit (SQK-LSK109) and the Ultra-Long DNA Sequencing Kit V14 (SQK-ULK114). Library preparation was performed according to the manufacturer’s protocol. Sequencing was carried out on the PromethION system using R10.4.1 flow cells (FLO-PRO114M) for ultra-long sequencing and R9.4.1 flow cells (FLO-PRO002) for conventional long-read sequencing. Basecalling was performed using the Dorado version 7.2 for ultra-long sequencing while guppy 3.2.8 and guppy 4.0.11 were used for basecalling of the conventional long-read sequencing.

For Hi-C analysis, we used 1–2 × 10^6^ cells per sample. The cells were fixed using a freshly prepared solution of 1% formaldehyde and 3 mM DSG in PBS. DSG was added first, and after 10 minutes, formaldehyde was introduced. The total fixation time was 20 minutes at room temperature, with continuous rotation on a rotator, following the manufacturer’s protocol. Hi-C DNA preparation was performed using the Dovetail Omni-C Kit, and the libraries were prepared with the Dovetail Library Module for Illumina 8Rx. To add sequencing indexes, we utilized the Dovetail Dual Index Primer Set #1 for Illumina 8Rx. The protocol followed was version 1.4 for mammalian samples. After chromatin digestion, enzyme concentrations were optimized to ensure that approximately 80% of the fragments fell within the 100–2500 bp range. Following proximity ligation, a size shift toward larger fragments was observed, a step we introduced that is not part of the original protocol. The final library DNA had an average fragment size of approximately 600 bp, which was then sequenced on an Illumina NovaSeq 6000.

Figure 1 illustrates basic statistics of the sequencing reads for Saudi and Japanese samples.

**Figure 1.**
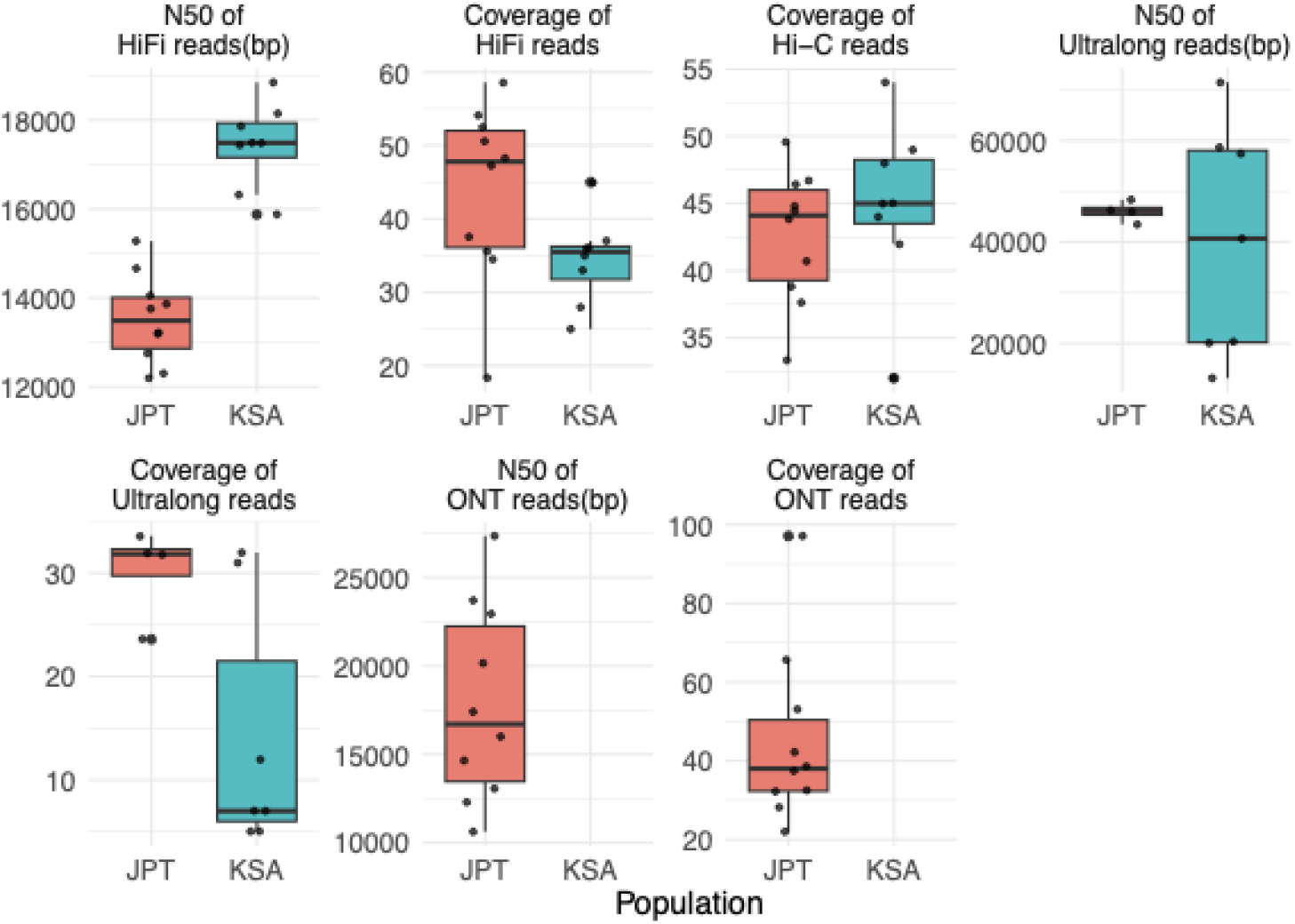
Overview of basic statistics of the sequencing reads from Japanese (JPT) and Saudi (KSA) samples.

### Genome assembly and assembly quality evaluation

For all samples, we primarily relied on a combination of PacBio HiFi reads, ONT reads, and Hi-C data to assemble the genomes and resolve haplotypes. Where available, we used ONT ultra-long (UL) reads for the assembly. We applied Hifiasm (UL) v0.19.0-r534 [13] in its Hi-C mode to generate haplotype-resolved assemblies. Hifiasm (UL) builds two string graphs based on HiFi and UL reads and then combines them. It uses the HiFi reads-based graphs as a backbone and merges them with UL reads-based graphs to make the assembly more complete. Furthermore, it uses Hi-C reads to resolve haplotypes and performs multiple rounds of graph cleaning. We generated all the assemblies with ONT UL or ONT reads except KSA007 for which we did not have ONT-based reads. We filtered out non-human contaminated sequences identified with Kraken2 [14] and ensured single mitochondrial assembly per maternal assembly.

### Genome annotation and identification of genes

We annotated the genome assemblies using Liftoff v1.6.1[15] with the T2T-CHM13 reference genome. Liftoff annotates a target genome by aligning reference gene sequences to the target genome using Minimap2. Minimap2 alignments are then parsed and fragmented alignments are combined using a directed acyclic graph. The graph represents gapless alignment blocks as nodes, connected by edges if they meet specific conditions.

This approach allows Liftoff to find the optimal mapping of reference genes to the target genome, even in cases of fragmented alignments. We counted mapped and unmapped genes and reported the results in the validation section.

Furthermore, we used Augustus v3.5.0 [16] and performed *de-novo* gene prediction on the assemblies. Then, we used Liftoff with Minimap2 to map the predicted genes to reference genomes GRCh38 and T2T-CHM13. We considered unmapped genes as novel genes and reported the statistics in the validation section.

### Pangenome Graph construction

We used Minigraph-Cactus pipeline v7.0.0 [17] to construct two distinct pangenome graphs for the Saudi Arabian and Japanese populations and one merged JaSaPaGe pangenome graph with samples from both populations, with GRCh38 and T2T-CHM13 as reference genomes. The Minigraph-Cactus pangenome pipeline is a five-step workflow for constructing a graph-based pangenome from multiple genome assemblies. The pipeline begins by constructing an initial structural variation (SV)-only graph using Minigraph, which iteratively adds variation from each assembly to a reference backbone without collapsing duplications. The graph is then augmented with smaller variants, and the input haplotypes are represented as paths. The pipeline includes steps for indexing, clipping, and post-processing the graph, including removing redundant sequences and nodes present in fewer than a specified number of haplotypes. The resulting graph is output in graphical fragment assembly (GFA) and VCF formats and can be used for mapping reads and calling variants. However, highly repetitive regions may require additional filtering to improve accuracy. We report basic statistics of variants present within the pangenome for JaSaPaGe, Japanese, Saudi Arabia and Human Pangenome Reference Consortium (HPRC) [1] pangenome graphs in Table 1. As evident from the statistics reported in Table 1, the number of detected variants in the merged JaSaPaGe pangenome graph is considerably higher than that of either the KSA or Japanese pangenome graphs.

**Table 1.** Pangenome graphs basic statistics. We report the number of samples from each population, without reference genomes included to provide shared coordinates.

| Pangenome | # samples<br>(without<br>reference) | Length | # SNPs | # MNPs | # Indels |
| --- | --- | --- | --- | --- | --- |
| HPRC | 47 | 3,234,968,488 | 20,758,652 | 446,253 | 7,359,580 |
| KSA | 9 | 3,159,673,318 | 9,475,991 | 322,642 | 2,196,775 |
| Japanese | 10 | 3,159,624,165 | 8,218,652 | 401,388 | 1,897,292 |
| JaSaPaGe | 19 | 3,177,502,842 | 11,395,931 | 495,539 | 2,549,282 |

### Short Read Variant Calling

To ensure the effectiveness of our approach, particularly in a clinical context where short-read sequencing is predominant, we employ a short-read variant calling workflow using the HPRC, Japanese, KSA, and JaSaPaGe pangenomes. This workflow is designed using the Common Workflow Language (CWL, v1.1). Our approach begins with data preparation, taking as input a reference genome and its index file, a variation graph, a haplotype index file, paired-end read files, and, optionally, specific genomic regions. We then align with VG Giraffe v1.54.0 [18], utilizing haplotype information to guide alignment. After that, we sort and index-aligned reads using Samtools v1.18, preparing them for statistical analysis and variant calling. Finally, we execute DeepVariant v1.6.0 [19] to detect variants. This step outputs a primary VCF file containing variant calls.

### Ethical approvals

The study with Saudi samples was approved by the Institutional Review Board (IRB) at King Fahad Medical City (KFMC) under approval number 22-037 and by the Institutional Bioethics Committee (IBEC) at King Abdullah University of Science and Technology (KAUST) under approval number 22IBEC023. The approval encompasses the recruitment of Saudi individuals with at least three generations of tribal roots from different regions of Saudi Arabia to provide blood samples. The protocol specifies that these samples will be sequenced using multiple sequencing technologies, processed with bioinformatics tools, and published without access restrictions.

Consent forms were available in both English and Arabic. Participants were recruited by Dr. Malak Abedalthagafi, the Principal Investigator (PI) for the approved IRB protocol at KFMC. Dr. Abedalthagafi personally obtained consent from the volunteers, who participated without any payment or compensation and were recruited from a research clinic. All consent forms were signed in the presence of two witnesses. There were no known professional or personal relationships between the authors and the participants, except for Dr. Abedalthagafi, who ensured that participants fully understood the risks and were competent to provide informed consent. As part of the consent process, all participants received genetic counseling as mandated by the IRB protocol.

None of the other authors were involved in the ethical approval processes at KFMC or KAUST. While Robert Hoehndorf was a member of the IBEC at KAUST at the time of approval, he did not participate in the review or discussion of the research protocol and was not present during its deliberations.

The Japanese sample was obtained from the JPT collection of the HapMap project, which comprises donors who have consented to unrestricted access to their genetic data.

## Data Records

The data described in this study are available in open access from public databases (DDBJ and NCBI Genbank). All data are available as a BioProject under the accession number PRJDB19680. This umbrella project includes separate projects for individual sequencing data, genome assemblies, and pangenome graphs.

## Technical Validation

### Validation of Genome Assemblies

We computed basic statistics and evaluated the quality of the assemblies using Merqury [20] with a k-mer size of 21. Figure 2 provides statistics on each of our assemblies. The number of contigs produced by the assemblies ranged from 118 to 2348 (mean: 405, median: 239); the N50 ranged from 10.046 to 106.755 MB (mean: 70.6, median: 80.9); the QV score computed by Merqury ranged from 56.2444 to 68.8636 (mean: 64.7, median: 65.3); the missing multi-copy gene (MMC) statistic ranged from 7.45% to 29.16% (mean: 12.4%, median: 10.7%); and the assembly length ranged from 2648.032 to 3141.660 MB (mean: 2981.5, median: 2977.5).

**Figure 2.**
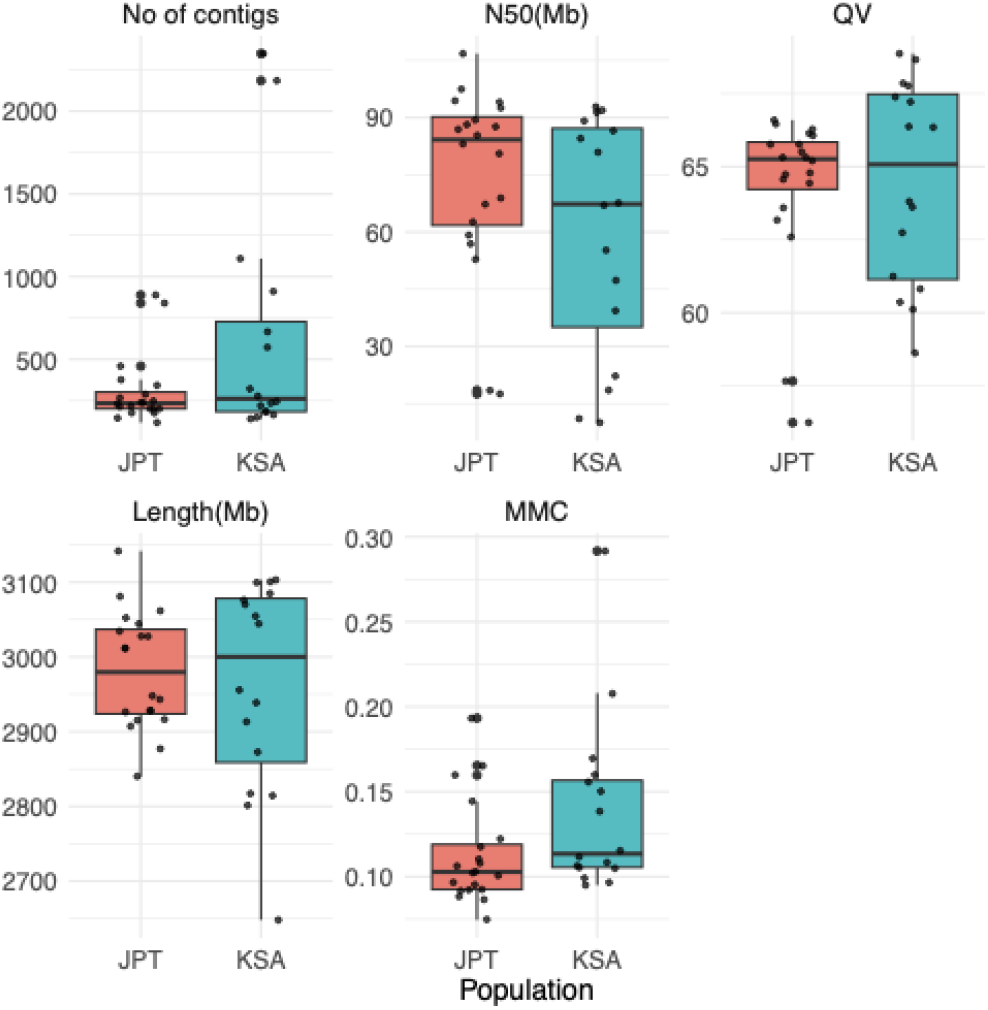
Overview of basic statistics of the genome assemblies from Japanese (JPT) and Saudi (KSA) samples.

Comparing the assemblies from Japanese and Saudi samples, Saudi samples showed more variability in assembly statistics. This can likely be attributed to differences in the quality and availability of sequencing data, as improved assembly quality belonged to the samples with more accurate long-read sequencing data, and lower assembly quality was observed for samples lacking nanopore sequencing data. There were no significant differences between the assemblies of the Japanese and Saudi samples with respect to the number of contigs, total length, N50, and QV score, but assemblies of the Japanese samples had a lower MMC than assemblies of the Saudi samples (p=0.048, Mann-Whitney U test).

As further validation of genome assembly, we performed a *de novo* gene annotation using Augustus [16] to identify potential novel genes and missing genes in the assemblies with respect to the T2T-CHM13v2.0 genome. Table 2 represents the summary of gene annotations. The number of protein-coding genes ranged from 16,257 to 19,101 (mean: 18,951, median: 19,096) with missing protein-coding genes ranging from 0 to 2,844 (mean: 173, median: 5); the total number of genes ranged from 55,137 to 60,880 (mean: 60,568, median: 60,862) with missing genes ranging from 88 to 5,831 (mean: 401, median: 106); and the number of novel genes ranged from 0 to 25 (mean: 2.1, median: 1). The assemblies of the Saudi samples had a significantly lower number of protein-coding genes and higher number of missing protein-coding genes, and we found no significant differences for other statistics.

**Table 2.**
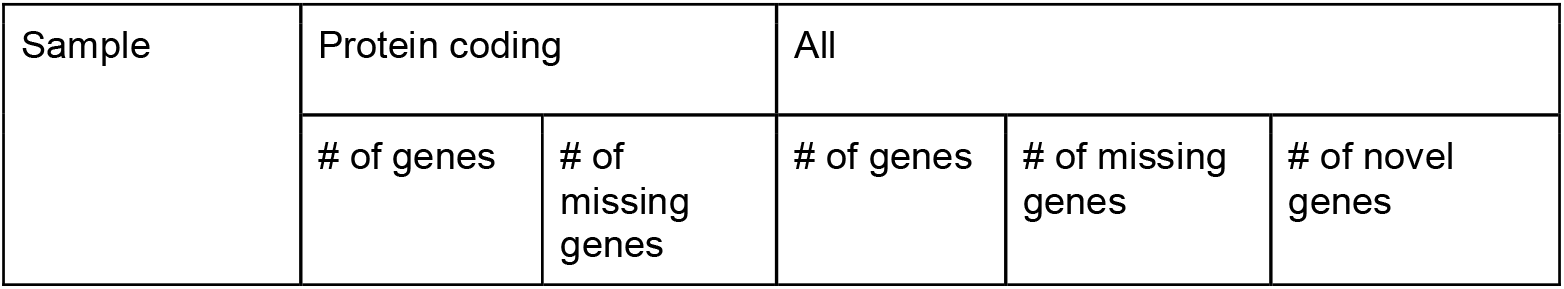
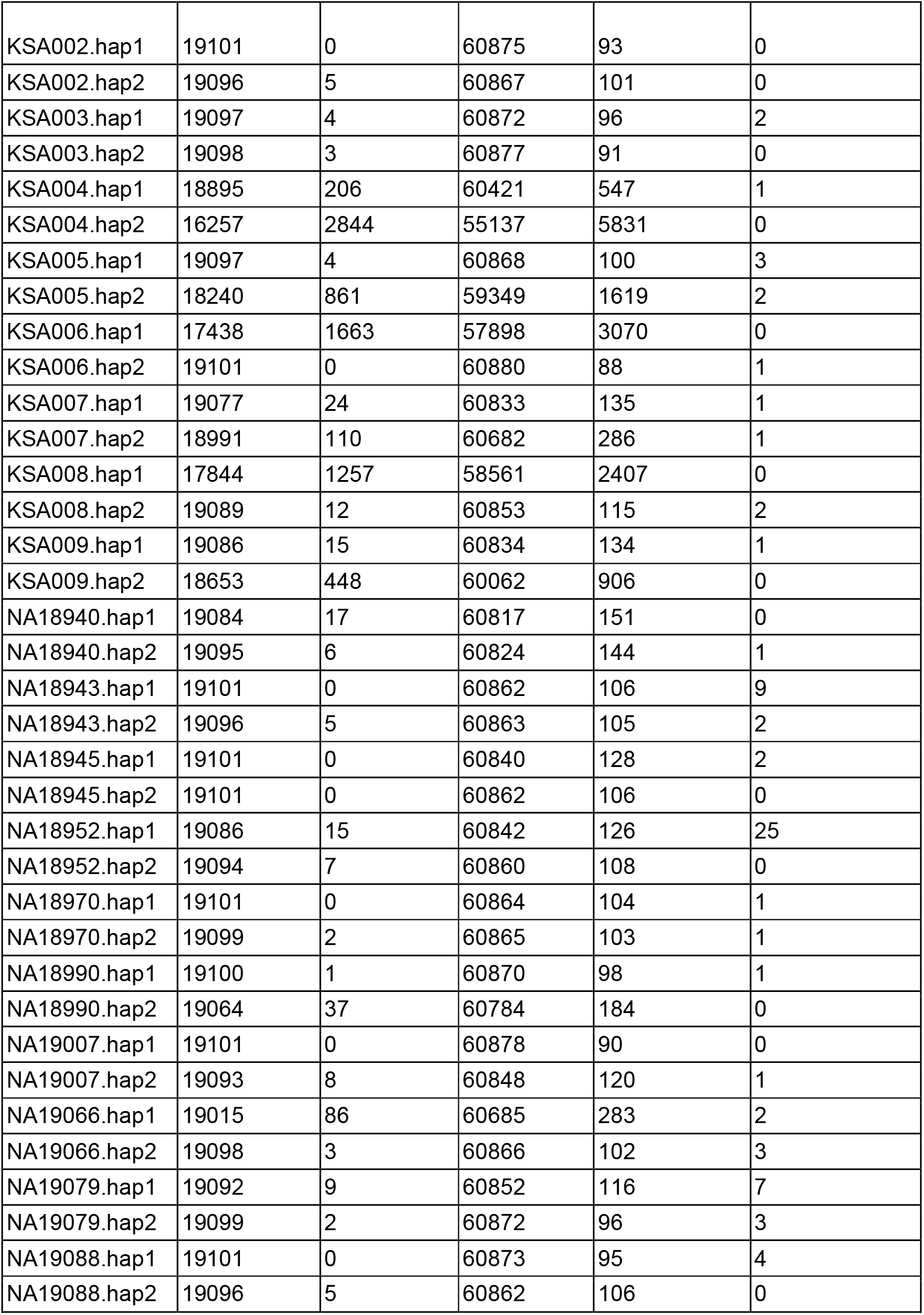
Annotation statistics.

| Sample | Protein coding |  | All |  |  |
| --- | --- | --- | --- | --- | --- |
|  | # of genes | # of missing genes | # of genes | # of missing genes | # of novel genes |
| KSA002.hap1 | 19101 | 0 | 60875 | 93 | 0 |
| KSA002.hap2 | 19096 | 5 | 60867 | 101 | 0 |
| KSA003.hap1 | 19097 | 4 | 60872 | 96 | 2 |
| KSA003.hap2 | 19098 | 3 | 60877 | 91 | 0 |
| KSA004.hap1 | 18895 | 206 | 60421 | 547 | 1 |
| KSA004.hap2 | 16257 | 2844 | 55137 | 5831 | 0 |
| KSA005.hap1 | 19097 | 4 | 60868 | 100 | 3 |
| KSA005.hap2 | 18240 | 861 | 59349 | 1619 | 2 |
| KSA006.hap1 | 17438 | 1663 | 57898 | 3070 | 0 |
| KSA006.hap2 | 19101 | 0 | 60880 | 88 | 1 |
| KSA007.hap1 | 19077 | 24 | 60833 | 135 | 1 |
| KSA007.hap2 | 18991 | 110 | 60682 | 286 | 1 |
| KSA008.hap1 | 17844 | 1257 | 58561 | 2407 | 0 |
| KSA008.hap2 | 19089 | 12 | 60853 | 115 | 2 |
| KSA009.hap1 | 19086 | 15 | 60834 | 134 | 1 |
| KSA009.hap2 | 18653 | 448 | 60062 | 906 | 0 |
| NA18940.hap1 | 19084 | 17 | 60817 | 151 | 0 |
| NA18940.hap2 | 19095 | 6 | 60824 | 144 | 1 |
| NA18943.hap1 | 19101 | 0 | 60862 | 106 | 9 |
| NA18943.hap2 | 19096 | 5 | 60863 | 105 | 2 |
| NA18945.hap1 | 19101 | 0 | 60840 | 128 | 2 |
| NA18945.hap2 | 19101 | 0 | 60862 | 106 | 0 |
| NA18952.hap1 | 19086 | 15 | 60842 | 126 | 25 |
| NA18952.hap2 | 19094 | 7 | 60860 | 108 | 0 |
| NA18970.hap1 | 19101 | 0 | 60864 | 104 | 1 |
| NA18970.hap2 | 19099 | 2 | 60865 | 103 | 1 |
| NA18990.hap1 | 19100 | 1 | 60870 | 98 | 1 |
| NA18990.hap2 | 19064 | 37 | 60784 | 184 | 0 |
| NA19007.hap1 | 19101 | 0 | 60878 | 90 | 0 |
| NA19007.hap2 | 19093 | 8 | 60848 | 120 | 1 |
| NA19066.hap1 | 19015 | 86 | 60685 | 283 | 2 |
| NA19066.hap2 | 19098 | 3 | 60866 | 102 | 3 |
| NA19079.hap1 | 19092 | 9 | 60852 | 116 | 7 |
| NA19079.hap2 | 19099 | 2 | 60872 | 96 | 3 |
| NA19088.hap1 | 19101 | 0 | 60873 | 95 | 4 |
| NA19088.hap2 | 19096 | 5 | 60862 | 106 | 0 |

### Validation of Pangenome Graphs using population-specific short read variant calling

We developed a variant-calling workflow to validate our population-specific pangenome graphs. We used Illumina short-read (150bp paired-end) sequencing data from three distinct sources: the NIST HG001 reference sample, three publicly available Saudi samples (SRR27002256, SRR29122519, SRR29095922), and three Japanese samples (NA18957, NA19064 and NA19086) from the high-coverage resequencing dataset of the International 1000 Genomes Project [21]. In addition to our pangenomes, we also used GRCh38, T2T-CHM13, and the HPRC pangenome graph as references for variant calling.

Tables 3, 4, and 5 show the statistics of variant calling for the Saudi samples, the Japanese samples, and HG001, respectively. We observed that using any pangenome graph as reference leads to a generally higher number of variants compared to using any linear reference genome (GRCh38 or T2T-CHM13). Furthermore, we found that variant calling generates more variants when the samples belong to the population of the pangenome; in particular, we identified more variants in Saudi samples when using the Saudi population-specific pangenome, and, similarly, we identified more variants in Japanese samples when using the Japanese population-specific pangenome. The results are as expected, as the population-specific pangenomes include population-specific variation which results in more reads being (correctly) aligned and, consequently, more variants being called. We also noticed a significant drop in variants being called when using T2T-CHM13 as reference, consistent with previous work [22] that observed the same phenomenon.

**Table 3.** Variant calling statistics for three Saudi samples, using Illumina reads aligned to GRCh38, T2T-CHM13, HPRC, KSA, Japanese, and JaSaPaGe.

| Reference | # Total number of variants | # SNPs | # Indels | #Homozygous variants | #Heterozygous variants |
| --- | --- | --- | --- | --- | --- |
| GRCh38 | 4,798,946 | 3,931,215 | 875,282 | 1,818,181 | 2,884,672 |
| T2T-CHM13 | 4,530,709 | 3,713,837 | 823,882 | 1,518,262 | 2,925,672 |
| HPRC | 4,918,192 | 4,006,467 | 914,641 | 1,883,276 | 2,928,903 |
| KSA | 4,929,121 | 4,016,643 | 914,954 | 1,890,125 | 2,933,112 |
| Japanese | 4,914,971 | 4,005,336 | 912,523 | 1,884,910 | 2,924,211 |
| JaSaPaGe | 4,924,157 | 4,011,559 | 915,018 | 1,886,809 | 2,931,466 |

**Table 4.** Variant calling statistics for the three Japanese samples, using Illumina reads aligned to GRCh38, T2T-CHM13, HPRC, KSA, Japanese, and JaSaPaGe.

| Reference | # Total number of variants | # SNPs | # Indels | #Homozygous variants | #Heterozygous variants |
| --- | --- | --- | --- | --- | --- |
| GRCh38 | 4,773,932 | 3,859,141 | 920,858 | 2,082,072 | 2,606,325 |
| T2T-CHM13 | 4,615,163 | 3,736,932 | 884,280 | 1,878,520 | 2,656,349 |
| HPRC | 4,874,741 | 3,945,970 | 932,478 | 2,176,336 | 2,608,881 |
| KSA | 4,877,008 | 3,949,027 | 931,614 | 2,180,319 | 2,607,461 |
| Japanese | 4,886,671 | 3,957,030 | 933,316 | 2,186,320 | 2,611,088 |
| JaSaPaGe | 4,880,980 | 3,951,546 | 932,978 | 2,181,795 | 2,609,804 |

**Table 5.** Variant calling statistics for the HG001 sample, using Illumina reads aligned to GRCh38, T2T-CHM13, HPRC, KSA, Japanese, and JaSaPaGe.

| Reference | # Total number of variants | # SNPs | # Indels | #Homozygous variants | #Heterozygous variants |
| --- | --- | --- | --- | --- | --- |
| GRCh38 | 4,772,087 | 3,859,738 | 918,507 | 1,894,665 | 2,791,773 |
| T2T-CHM13 | 4,478,813 | 3,626,338 | 858,191 | 1,551,671 | 2,851,517 |
| HPRC | 4,868,841 | 3,943,065 | 929,514 | 1,982,634 | 2,796,852 |
| KSA | 4,876,386 | 3,950,714 | 929,353 | 1,990,503 | 2,796,673 |
| Japanese | 4,870,049 | 3,945,318 | 928,474 | 1,986,476 | 2,794,423 |
| JaSaPaGe | 4,869,064 | 3,943,503 | 929,114 | 1,983,690 | 2,796,126 |

In addition to validation of the generated population-specific pangenomes, we compared variant calling results of short-read sequencing data for the HG001 genome to the truth set of variants provided by NIST. Tables 5 and 6 show the performance evaluation for calling SNVs and indels, respectively. We observed that the variant calling performance is comparable when either our pangenome graphs (KSA, Japanese, and JaSaPaGe) or HPRC pangenome graph are used as reference, and it is much improved than when GRCh38 is used as reference.

**Table 5.**
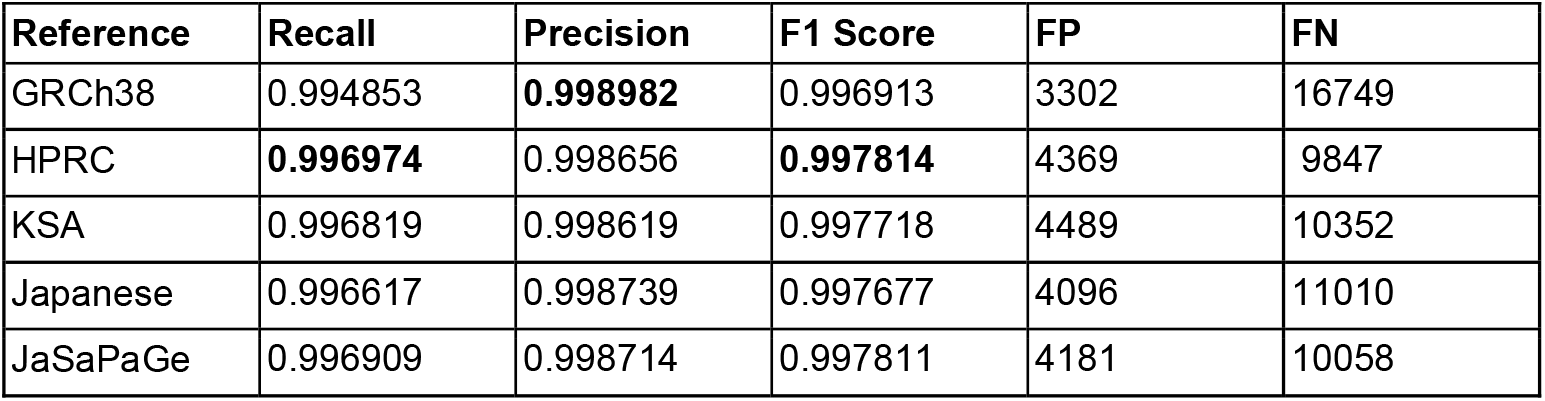
SNV performance statistics for the HG001 sample, using Illumina reads aligned to GRCh38, HPRC, KSA, Japanese, and JaSaPaGe. Bold font indicates best performance. FP indicates false positives and FN false negatives.

| Reference | Recall | Precision | F1 Score | FP | FN |
| --- | --- | --- | --- | --- | --- |
| GRCh38 | 0.994853 | **0.998982** | 0.996913 | 3302 | 16749 |
| HPRC | **0.996974** | 0.998656 | **0.997814** | 4369 | 9847 |
| KSA | 0.996819 | 0.998619 | 0.997718 | 4489 | 10352 |
| Japanese | 0.996617 | 0.998739 | 0.997677 | 4096 | 11010 |
| JaSaPaGe | 0.996909 | 0.998714 | 0.997811 | 4181 | 10058 |

**Table 6.** INDEL performance statistics for the HG001 sample, using Illumina reads aligned to GRCh38, HPRC, KSA, Japanese, JaSaPaGe. Bold font indicates best performance. FP indicates false positives and FN false negatives.

| Reference | Recall | Precision | F1 Score | FP | FN |
| --- | --- | --- | --- | --- | --- |
| GRCh38 | 0.991815 | **0.995341** | 0.993575 | 2255 | 3828 |
| HPRC | 0.992863 | 0.994684 | 0.993773 | 2578 | 3338 |
| KSA | 0.992842 | 0.994786 | 0.993813 | 2528 | 3348 |
| Japanese | 0.992705 | 0.994802 | 0.993752 | 2520 | 3412 |
| JaSaPaGe | **0.99291** | 0.99477 | **0.993839** | 2536 | 3316 |

### Evaluation of FAIR principles

We evaluated the pangenome resources provided in this study against the FAIR principles [11]. The pangenome is Findable in the DDBJ database [23] under the globally unique identifier PRJDB19680, with comprehensive metadata adhering to DDBJ’s standardized requirements for genomic data submission. The resource is Accessible through DDBJ’s established protocols, which are open, free, and universally implementable via standard web interfaces and APIs.

For Interoperability, the pangenomes are provided in the standard GFA format [24], with explicit links to the constituent genome assemblies and their corresponding primary sequencing data accessions. To ensure Reusability, we have documented the complete analysis workflow, including all computational methods used to generate the pangenome, and have made the data publicly available. All associated metadata and data formats comply with community standards for pangenome representation and data exchange.

## Code Availability

Source code and additional resources are available at https://github.com/JaSaPaGe/pangenome-cwl.

## Acknowledgments

We acknowledge support from the KAUST Supercomputing Laboratory and the KAUST Bioscience Core Lab. Part of the computational analysis was performed on the NIG supercomputer at the ROIS National Institute of Genetics.

We extend our gratitude to the organizers and participants of Biohackathon 2023 and 2024, as well as to DBCLS, whose support and collaborative environment provided the inspiration and facilitated the initial discussions for this research.

We also thank the eight anonymous blood donors for their invaluable contributions, providing samples KSA002 to KSA009, which were essential for this research.

This work was supported by King Abdullah University of Science and Technology (KAUST) Office of Sponsored Research (OSR) (REI/1/5659-01-01) and by funding from King Abdullah University of Science and Technology – KAUST Center of Excellence for Smart Health (KCSH),(5932) to R.H. Y.K. and M.N. are supported by the Database Integration

Coordination Program (DICP) of the National Bioscience Database Center NBDC of Japan Science and Technology Agency (JPMJND2302). Y.O. was supported by Grants-in-Aid for Scientific Research from the Ministry of Education, Culture, Sports, Science, and Technology (JP18H05527 and JP24H02323); the Medical Research Center Initiative for High Depth Omics, Kyushu University, Japan; MEXT Promotion of Development of a Joint Usage/Research System Project: the Cooperative Research Project Program ; MEXT Promotion of Development of a Joint Usage/Research System Project: Coalition of Universities for Research Excellence Program (CURE) (JPMXP1323015486). M.S.A is supported by funding from King Salman Center for Disability Research, grant R-20190016. The infrastructure of Omics Science Center Secure Information Analysis System, Medical Institute of Bioregulation at Kyushu University provides the (part of) computational resource (https://sis.bioreg.kyushu-u.ac.jp/). This work was supported in part by the MEXT Cooperative Research Project Program, Medical Research Center Initiative for High Depth Omics, and CURE:JPMXP1323015486 for MIB, Kyushu University. This work was partly performed in the Cooperative Research Project Program of the Medical Institute of Bioregulation, Kyushu University. Masao Nagasaki received grants from the Japan Agency for Medical Research and Development (AMED) (Grant Numbers JP21wm0425009, JP22fk0210111, JP22tm0424222, JP23ek0109675, JP23ek0109672, JP23fk0210138, JP23ek0210194, JP24gm2010001) and JST NBDC Grant Number JPMJND2302, and JSPS KAKENHI Grant Number JP21H02681. This work was partially supported by the “Joint Usage/Research Center for Interdisciplinary Large-scale Information Infrastructures” and “High Performance Computing Infrastructure” in Japan (Project ID: jh230016, and jh240015).

## Author Information

### Authors and Affiliations

**Computer, Electrical and Mathematical Sciences & Engineering (CEMSE) Division, King Abdullah University of Science and Technology, King Abdullah University of Science and Technology, 4700 KAUST, Thuwal, Saudi Arabia** Maxat Kulmanov, Marwa Abdelhakim, Robert Hoehndorf

**KAUST Center of Excellence for Smart Health (KCSH), King Abdullah University of Science and Technology, 4700 KAUST, Thuwal, Saudi Arabia**

Maxat Kulmanov, Yang Liu, Marwa Abdelhakim, Rund Tawfiq, Robert Hoehndorf

**KAUST Center of Excellence for Generative AI, King Abdullah University of Science and Technology, 4700 KAUST, Thuwal, Saudi Arabia**

Maxat Kulmanov, Rund Tawfiq, Robert Hoehndorf

**SDAIA–KAUST Center of Excellence in Data Science and Artificial Intelligence, King Abdullah University of Science and Technology, 4700 KAUST, Thuwal, Saudi Arabia** Maxat Kulmanov, Robert Hoehndorf

**Genome Medical Science Project, Research Institute, National Center for Global Health and Medicine, Shinjuku-ku, Tokyo 162-8655, Japan**

Saeideh Ashouri, Katsushi Tokunaga, Yosuke Kawai

**Biological and Environmental Sciences & Engineering (BESE) Division, King Abdullah University of Science and Technology, King Abdullah University of Science and Technology, 4700 KAUST, Thuwal, Saudi Arabia**

Yang Liu, Rund Tawfiq, Robert Hoehndorf

**Genomics and Precision Medicine Department, King Fahad Medical City, Riyadh, Saudi Arabia**

Ebtehal Alsolm

**Division of Biomedical Information Analysis, Medical Research Center for High Depth Omics, Medical Institute of Bioregulation, Kyushu University**,, **Fukuoka 812-0054 Japan**

Masao Nagasaki

**Center for Genomic Medicine, Graduate School of Medicine, Kyoto University, Kyoto 606-8507 Japan**

Masao Nagasaki

**Division of Transcriptomics, Medical Institute of Bioregulation, Kyushu University**,, **Fukuoka 812-0054 Japan**

Yasuyuki Ohkawa

**Department of Computational Biology and Medical Sciences, Graduate School of Frontier Sciences, The University of Tokyo, Chiba, 277-8561 Japan**

Yutaka Suzuki

**Database Center for Life Science, Joint Support-Center for Data Science Research, Research Organization of Information and Systems, Chiba, 277-0871 Japan** Toshiaki Katayama

**Department of Pathology and Laboratory Medicine, Emory School of Medicine, Atlanta, GA, USA**

Malak S Abedalthagafi

**King Salman Center for Disability Research, Riyadh, Saudi Arabia**

Malak S Abedalthagafi

**Bioinformation and DDBJ Center, Research Organization of Information and Systems, National Institute of Genetics, Shizuoka, 411-8540 Japan**

Yosuke Kawai

## Contributions

Y.K., K.T., T.K., R.H., and M.S.A. designed this study and managed the progress of the project. M.S.A and E.A. conducted the recruitment and consent. M.N., Y.O., Y.S., M.Abd., R.T., and E.A. conducted the sample processing. K.T., M.N., Y.O., Y.S., and M.Abd. conducted the library preparation and sequencing. Y.K., S.A., T.K., M.K., Y.L., conducted the data processing and bioinformatics analysis.

## Corresponding authors

Correspondence to Malak S. Abedalthagafi, Robert Hoehndorf, and Yosuke Kawai.

